# AHA1 regulates Aβ production via modulation of APP expression and γ-secretase assembly

**DOI:** 10.1101/2025.11.03.686106

**Authors:** Arshad Ali Noorani, Sadequl Islam, Kevin Catalfano, Heather M. Wilkins, Brian S. J. Blagg, Kun Zou, Michael S. Wolfe

**Author notes:** **To whom correspondence should be addressed:** Arshad Ali Noorani Department of Pharmacology & Immunology, Medical University of South Carolina, Charleston, SC 29425, USA. and Michael S. Wolfe Department of Medicinal Chemistry, University of Kansas, Lawrence, KS 66045, USA.

## Abstract

Deposition of amyloid β-protein (Aβ) is a hallmark of Alzheimer’s disease (AD), produced by γ-secretase–mediated cleavage of amyloid precursor protein (APP). The 90-kDa heat shock protein (Hsp90) co-chaperone, activator of Hsp90 ATPase homolog 1 (AHA1), is known to promote the accumulation of toxic tau species; however, its effects on Aβ production remain unclear. Here, we show that knockdown of endogenous AHA1 decreases Aβ generation and reduces APP and γ-secretase components, whereas AHA1 overexpression elevates Aβ production and the expression of these proteins. The AHA1-E67K mutant, which has impaired Hsp90 binding, lowers Aβ production and the levels of APP and γ-secretase components compared with wild-type AHA1. AHA1 associates with APP and immature γ-secretase components, including anterior pharynx-defective phenotype 1 (APH1), indicating its role in APP proteolysis and Aβ production. Disruption of the AHA1/Hsp90 complex—through AHA1 knockdown, the E67K mutant, or a small-molecule inhibitor—reduces γ-secretase assembly. Familial AD mutations in APP-C99 and presenilin-1 (PS1) increase AHA1, Hsp90, APP, and APH1 expression, enhancing Aβ production. Importantly, AHA1 knockdown decreases abnormal Aβ generation and C99, PS1-CTF, and APH1 levels in mutant APP cells, while AHA1 overexpression enhances Aβ production in PS1 mutant cells. Collectively, these findings reveal that AHA1 regulates Aβ production by modulating APP expression and γ-secretase assembly, establishing AHA1 as a potential target for therapeutic intervention in AD.

## Introduction

Alzheimer’s disease (AD) is characterized by the formation of amyloid plaques and neurofibrillary tangles (NFTs) made up of aggregated β-amyloid (Aβ) and tau, respectively (1). Aβ peptides are products of the sequential processing of amyloid precursor protein (APP) by the β-amyloid cleaving enzyme 1 (BACE1) and the γ-secretase complex (2), comprised of presenilin (PS), nicastrin (NCT), anterior pharynx-defective phenotype 1 (APH1), and presenilin enhancer 2 (PEN-2) (3-5). AD is broadly categorized based on its onset and genetic factors into Sporadic Alzheimer’s Disease (SAD) and Familial Alzheimer’s Disease (FAD). Autosomal dominant mutations in APP, PS1, and PS2 are associated with early-onset FAD and result in increased molar ratios of Aβ42/Aβ40, which is uncommon compared to late-onset SAD (6-9). The exact cause of SAD is less clear and is believed to result from a complex interplay of genetic, environmental, and lifestyle factors (10). Both SAD and FAD share common clinical features, such as progressive memory loss and cognitive decline, as well as pathological hallmarks, including accumulation of Aβ plaques and tau protein tangles in the brain (11, 12). Nevertheless, the precise mechanisms and contributions of Aβ to disease pathogenesis and progression continue to be subjects of active investigation (13, 14).

Molecular chaperones and stress-response proteins regulate the folding, stability, and clearance of proteins and are particularly important in preventing dysfunction and aggregation of mutated and misfolded proteins (15). Many studies have demonstrated that heat shock protein 90 (Hsp90) is the well-known molecular chaperone protein that assists in the proper folding of other proteins, stabilizes them under stress, and aids in their degradation (16, 17). Recently, we reported that high temperature regulates Aβ production by modulating γ-secretase complex formation through the interaction of Hsp90 to the APH1/NCT partially assembled γ-secretase complex (18). Previous findings have indicated that Hsp90 can influence both Aβ production and tau accumulation through its direct association with γ-secretase complexes and tau protein (19-21). However, for Hsp90 to function properly, co-chaperones regulate its enzymatic (ATPase) and folding activities and guide specific proteins to interact with Hsp90, as it does not operate independently (22). One such Hsp90 co-chaperone is the activator of Hsp90 ATPase homolog 1 (AHA1), which stimulates the ATPase activity of Hsp90. Notably, AHA1 is the only known activator of Hsp90 ATPase activity in mammals (23, 24). This small 38-kDa co-chaperone binds to the N-terminal and middle domains of Hsp90, inducing a partially closed conformation that dramatically accelerates the ATPase cycle’s progression (25, 26). Apart from its role in stimulating the ATPase activity of Hsp90, AHA1 also functions in the late stage of the Hsp90 chaperone cycle. In addition, AHA1 can act independently of Hsp90 as an autonomous chaperone to prevent the aggregation of stressed or misfolded proteins (27-30). Previous research has indicated that individuals with AD have increased levels of AHA1/Hsp90 in their brains when compared to age-matched controls (31). In addition, AHA1 has been found to increase the production of harmful tau aggregates (32, 33). On the other hand, it has been observed that AHA1 regulates the expression of Dicer1 protein, which plays a key role in the development of cancer (34). Another report has suggested that AHA1 may play an oncogenic role in osteosarcoma by increasing the levels of isocitrate dehydrogenase 1 (IDH1) (35). Moreover, AHA1 has been found to increase the misfolding of cystic fibrosis transmembrane conductance regulator, including the Δ508 mutant that is commonly seen in cystic fibrosis patients (36). However, the role of AHA1 in regulating the other proteins is largely unknown. Therefore, identifying the proteins that AHA1 interacts with in the Hsp90 network will help us understand AHA1’s role in biological processes regulated by Hsp90.

In this study, we hypothesized that AHA1 may regulate the production of Aβ independently of Hsp90 interaction or by modifying Hsp90 activity. We demonstrated that AHA1 can regulate the levels of APP independently of Hsp90 interaction and the formation of the γ-secretase complex through a pathway that depends on Hsp90 interaction, ultimately affecting Aβ production. Our findings show that AHA1 is associated with APP and the components of the immature γ-secretase complex, APH1, highlighting the role of AHA1 in modulating APP and γ-secretase assembly. Additionally, we found a novel link between the upregulation of AHA1 function and FAD mutations, connecting the mechanism associated with loss of function in FAD mutations and abnormal Aβ production. For the first time, our findings indicate a novel mechanism by which AHA1 is involved in AD pathogenesis through the regulation of Aβ production.

## Results

### AHA1 regulates the production of Aβ and the levels of APP independently of its interaction with Hsp90

Because AHA1 can affect the accumulation of toxic tau species(32), we investigated whether altering AHA1 expression can affect Aβ production. To determine the role of AHA1 in Aβ production, we performed the knockdown of AHA1 by transfecting with siRNA in human embryonic kidney 293T (HEK293T) cells, stably overexpressing human APP695. Interestingly, we found that the levels of Aβ40 and Aβ42 were significantly decreased in the culture medium of AHA1 siRNA transfected cells compared to control siRNA-transfected cells (Fig. 1A). We also examined whether AHA1 overexpression enhances Aβ production in HEK293T cells. We co-transfected HEK293T cells with human APP695 and either wild-type (WT) AHA1 or AHA1 E67K (a mutant that cannot bind to Hsp90) or an empty vector (EV) control. We found that WT AHA1 significantly increased the levels of secreted Aβ40 and Aβ42 compared to EV control, whereas AHA1 mutants AHA1 E67K enhanced Aβ40 and Aβ42 secretion to a lesser degree than WT AHA1 (Fig. 1B). These results indicate that AHA1 positively regulates Aβ40 and Aβ42 production in this system.

**Figure 1.**
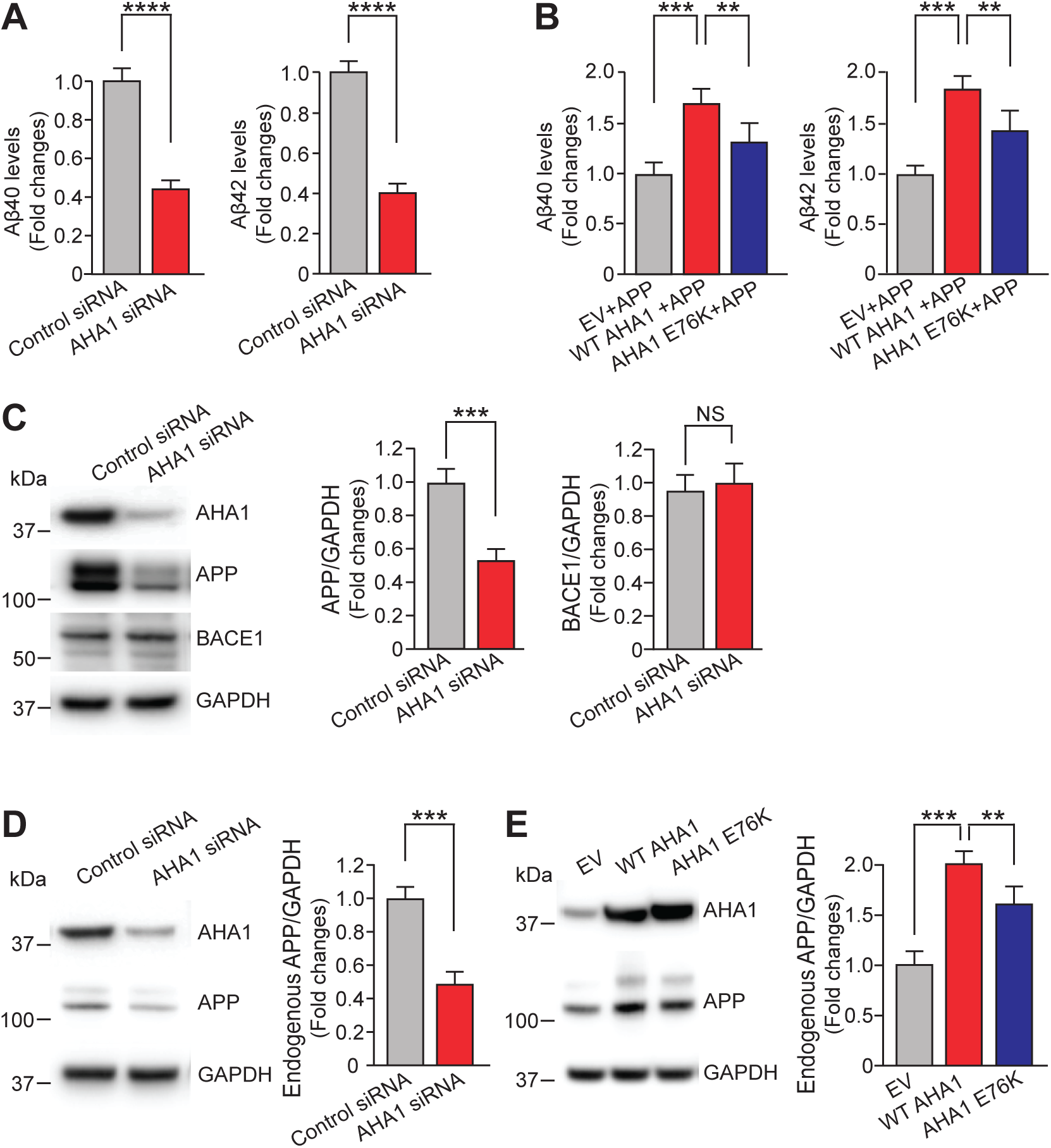
AHA1 controls Aβ production and APP levels independently of its interaction with Hsp90. **(A)** The amount of secreted Aβ40 and Aβ42 levels in the media of HEK-APP cells transfected with AHA1 siRNA or Control siRNA were measured with an Aβ ELISA kit. **(B)** The amount of secreted Aβ40 and Aβ42 levels in the media of HEK293T cells transiently co-transfected with APP695, and either WT AHA1 or AHA1 E67K or EV were measured with an Aβ ELISA kit. **(C)** The expression levels of AHA1, full-length APP, and BACE1 in HEK-APP cells after the knockdown of AHA1 were detected with Western blotting. **(D)** The expression levels of AHA1 and full-length endogenous APP in HEK293T cells after knockdown of AHA1 were detected with Western blotting. **(E)** The expression levels of AHA1 and full-length endogenous APP in HEK293T cells after overexpression of WT AHA or AHA1 E67K mutant were determined using Western blotting. Data are the mean ± S.E. (error bars) from three independent experiments. **, p < 0.01; ***, p < 0.001; ****P < 0.0001; as determined with Student’s *t* test and one-way ANOVA with Tukey’s multiple-comparison test.

To gain mechanistic insight into how AHA1 affects Aβ production, we investigated the cellular levels of APP and BACE1 in HEK-APP cells where AHA1 expression was knocked down using siRNA. Notably, we found that the cellular levels of APP were dramatically reduced; however, the BACE1 levels remained unchanged in AHA1 knocked-down cells compared to control siRNA-transfected cells (Fig. 1C). This finding implies that the reduction of AHA1 in cells leads to a decrease in exogenous APP levels.

Furthermore, the AHA1 knock-down also reduced endogenous APP expression in HEK293T cells (Fig. 1D). In addition, transfection of WT AHA1 and mutant AHA1 E67K in HEK293T cells resulted in an increase in endogenous APP levels compared to EV control (Fig. 1E). Taken together, these results suggest that AHA1 can regulate Aβ production by modulating APP expression levels irrespective of its interaction with Hsp90.

### AHA1 modulates the **γ**-secretase components and its activity through a pathway that is dependent on Hsp90 interaction

AHA1 is known to stimulate the ATPase activity of Hsp90, thereby promoting the folding of its client proteins (23). While reducing Hsp90 levels decreased γ-secretase components (18), it is unknown whether altering the Hsp90 activity by AHA1 affects γ-secretase levels. Interestingly, we found that knock-down of AHA1 by using AHA1 siRNA significantly reduced the levels of C-terminal fragment of presenilin 1 (PS1-CTF), NCT, APH1, and PEN-2 γ-secretase components compared to control siRNA (Fig. 2A). In contrast, transient co-transfection of WT AHA1 and APP (AHA1+APP) in HEK293T cells remarkably increased the levels of PS1-CTF, NCT, APH1, and PEN-2 compared to EV control (Fig. 2B). These findings suggest that AHA1 modulates the expression of γ-secretase molecules by altering Hsp90 activity.

**Figure 2.**
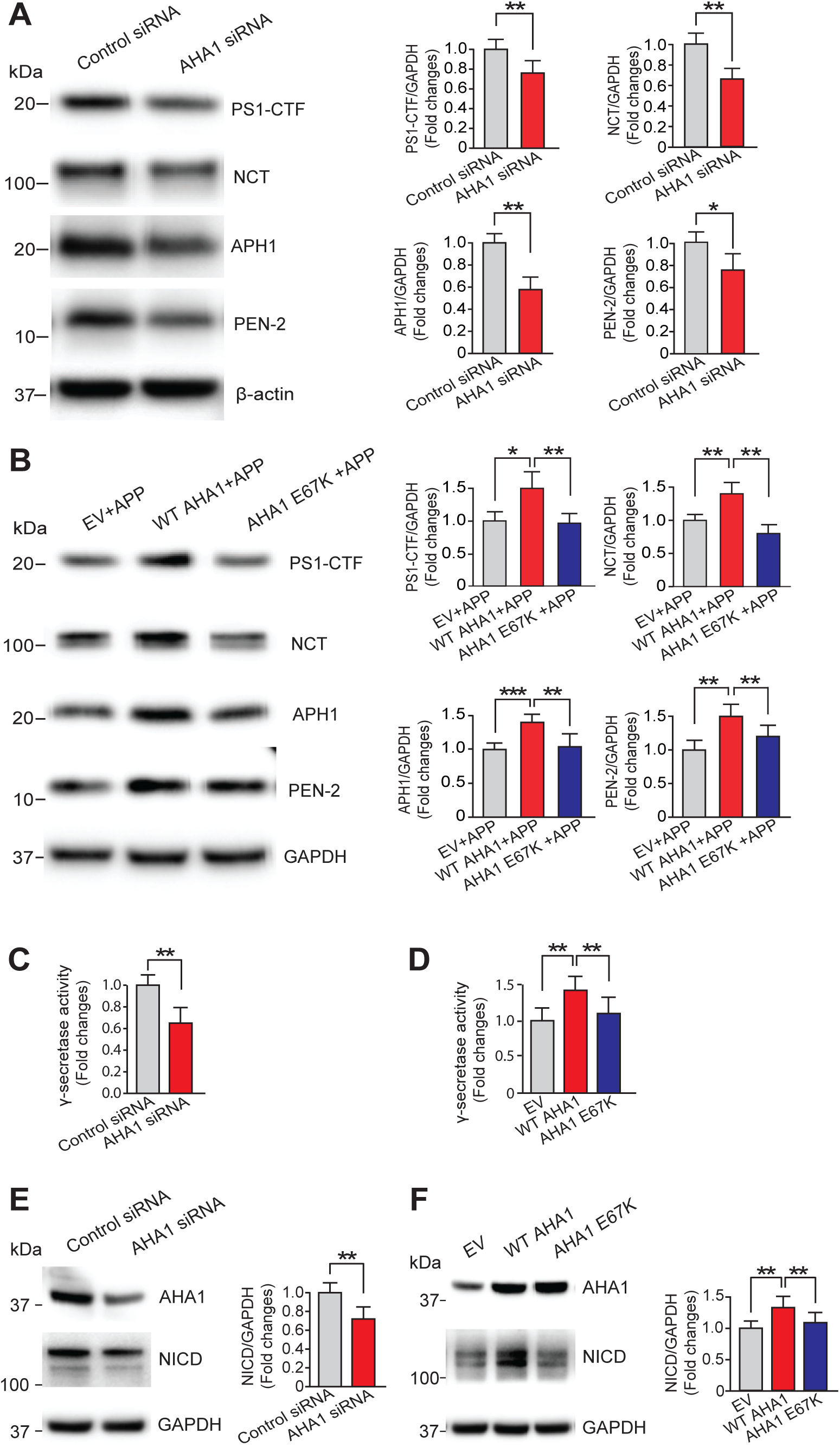
AHA1 modulates the components and activity of γ-secretase through an Hsp90-dependent pathway. **(A)** The expression levels of PS1-CTF, NCT, APH1, and PEN-2 in HEK-APP cells after the knockdown of AHA1 were detected with Western blotting. **(B)** The expression levels of PS1-CTF, NCT, APH1, and PEN-2 in HEK293T cells after overexpression of WT AHA1 or AHA1 E67K mutant were detected with Western blotting. **(C)** γ-secretase activity was detected with a fluorogenic substrate after the knockdown of AHA1 in HEK293T cells. **(D)** γ-secretase activity was detected with a fluorogenic substrate after overexpression of WT AHA1 or AHA1 E67K mutant in HEK293T cells. **(E)** The expression levels of AHA1 and NICD in WT MEF cells after the knockdown of AHA1 were detected with Western blotting. **(F)** The expression levels of AHA1 and NICD in WT MEF cells after overexpression of WT AHA1 or AHA1 E67K mutant were detected with Western blotting. Data are the mean ± S.E. (error bars) from three independent experiments. *, p < 0.05; **, p < 0.01; ***, p < 0.001 as determined with Student’s *t* test and one-way ANOVA with Tukey’s multiple-comparison test.

To investigate whether this modulation depends on the interaction between AHA1 and Hsp90, we co-transfected Hsp90-binding-defective mutant AHA1 E67K and APP in HEK293T cells. As expected, the mutant AHA1 E67K+APP overexpression decreased the levels of γ-secretase components compared to WT AHA1+APP overexpression or with levels equivalent to the EV+APP overexpression (Fig. 2B). Consistent with this finding, we also observed a significant decrease in the complex formation between Hsp90 and AHA1 in HEK293T cells upon treatment with the AHA1/Hsp90 small molecule disruptor ND-AHA1-46 (Fig. S1A, B). ND-AHA1-46, was developed by our group to disrupt interactions between AHA1 and Hsp90. Previous research has shown that ND-AHA1-46 can reduce P301L tau aggregation *in vitro* (37). Furthermore, we also observed the impact of blocking AHA1/Hsp90 complex formation in HEK-APP cells for 24 h using ND-AHA1-46 on the expression levels of γ-secretase components. Blockade of AHA1/Hsp90 complex formation using ND-AHA1-46 for 24h reduced the levels of N-terminal fragment of presenilin 1 (PS1-NTF), NCT, APH1, and PEN-2 γ-secretase molecules compared to DMSO control (Fig. S2A). However, the cellular levels of APP were significantly increased in ND-AHA1-46-treated cells (Fig. S2A). In addition, using blue-native polyacrylamide gel electrophoresis (BN-PAGE), the formation of PS1-NTF, NCT, and APH1 γ-secretase complex was also remarkably downregulated in ND-AHA1-46-treated cells (Fig. S2B). Taken together, these findings suggest that modulation of the expression of γ-secretase molecules by AHA1 depends on the AHA1/Hsp90-associated mechanism.

To identify the role of AHA1 on γ-secretase assembly in the absence of PS, we observed the precomplex formation in PS1/2 double-KO (PS-DKO) mouse embryonic fibroblasts (MEFs) at ∼240 kDa. We found that blockade of AHA1/Hsp90 interaction using ND-AHA1-46 for 24 h significantly reduced the precomplex formation compared to DMSO control (Fig. S2C). This result suggests that AHA1/Hsp90 interaction is also crucial for γ-secretase assembly irrespective of the PS-mediated γ-secretase assembly.

Furthermore, we also found that γ-secretase activity decreased in the AHA1 knockdown HEK293T cells by an *in vitro* γ-secretase activity assay (Fig. 2C). Overexpression of WT AHA1 in HEK293T cells increased the γ-secretase activity compared to EV control (Fig. 2D). AHA1 E67K decreased the levels of γ-secretase activity compared to WT AHA1 (Fig. 2D). Overexpression of AHA1 E67K did not change γ-secretase activity levels compared to EV control. These results indicate that AHA1 regulates the γ-secretase activity in an Hsp90-dependent manner.

Because high temperature increases the levels of Hsp90 and Notch1 intracellular domain (NICD) in mouse brains (18), we further investigated whether AHA1 can also affect the cleavage of other substrates of γ-secretase. We used WT MEF cells because HEK293T cells express trace endogenous NICD. Notably, we found that knock-down of AHA1 significantly decreased the NICD levels in WT MEF cells (Fig. 2E). Overexpression of WT AHA1 increased the NICD levels in WT MEF cells (Fig. 2F). However, mutant AHA1 E67K decreased the levels of NICD compared to WT AHA1 (Fig. 2F). Overexpression of AHA1 E67K did not change NICD levels compared to EV control (Fig. 2F). These findings indicate that AHA1 can play a role in regulating the γ-secretase mediated cleavage of notch. Collectively, these results suggest that AHA1 regulates the components and activity of γ-secretase through a pathway that depends on Hsp90 interaction.

### AHA1 associates with APP and APH1

To investigate whether AHA1 is involved in the regulation of APP and the formation of the γ-secretase complex, we performed immunoprecipitation (IP) experiments using HEK293T cells. We found that AHA1 is bound to APP (Fig. 3A) and APH-1 but not the other γ-secretase components, PS1-NTF, NCT, or PEN-2 (Fig. 3B). These results suggest that AHA1 is involved in the regulation of APP and the formation of the γ-secretase complex. However, AHA1 did not interact with BACE1 (Fig. 3B), suggesting that AHA1 regulates Aβ production through interacting with APP and γ-secretase components without hampering β-secretase activity. To confirm further that AHA1 regulates γ-secretase by a pathway dependent on Hsp90 interaction, we overexpressed the WT AHA1 and AHA1 E67K mutant in HEK-293T cells and performed IP experiments. As reported previously(32), we found that AHA1 E67K showed a weaker interaction with Hsp90 than WT AHA1 (Fig. 3C). Interestingly, AHA1 E67K interacts with less APH1 compared with WT AHA1 (Fig. 3C). These results suggest that the AHA1/Hsp90 complex plays a crucial role in regulating γ-secretase assembly. In addition, confocal microscopy studies demonstrated that colocalization between AHA1 and APP or APH1 (Fig. 3D) is consistent with the findings in the IP studies. These results suggest that AHA1 may promote the assembly of the γ-secretase, possibly first by forming the precomplex through its interaction with APH1 and modulating Aβ production by interacting with APP and APH1.

**Figure 3.**
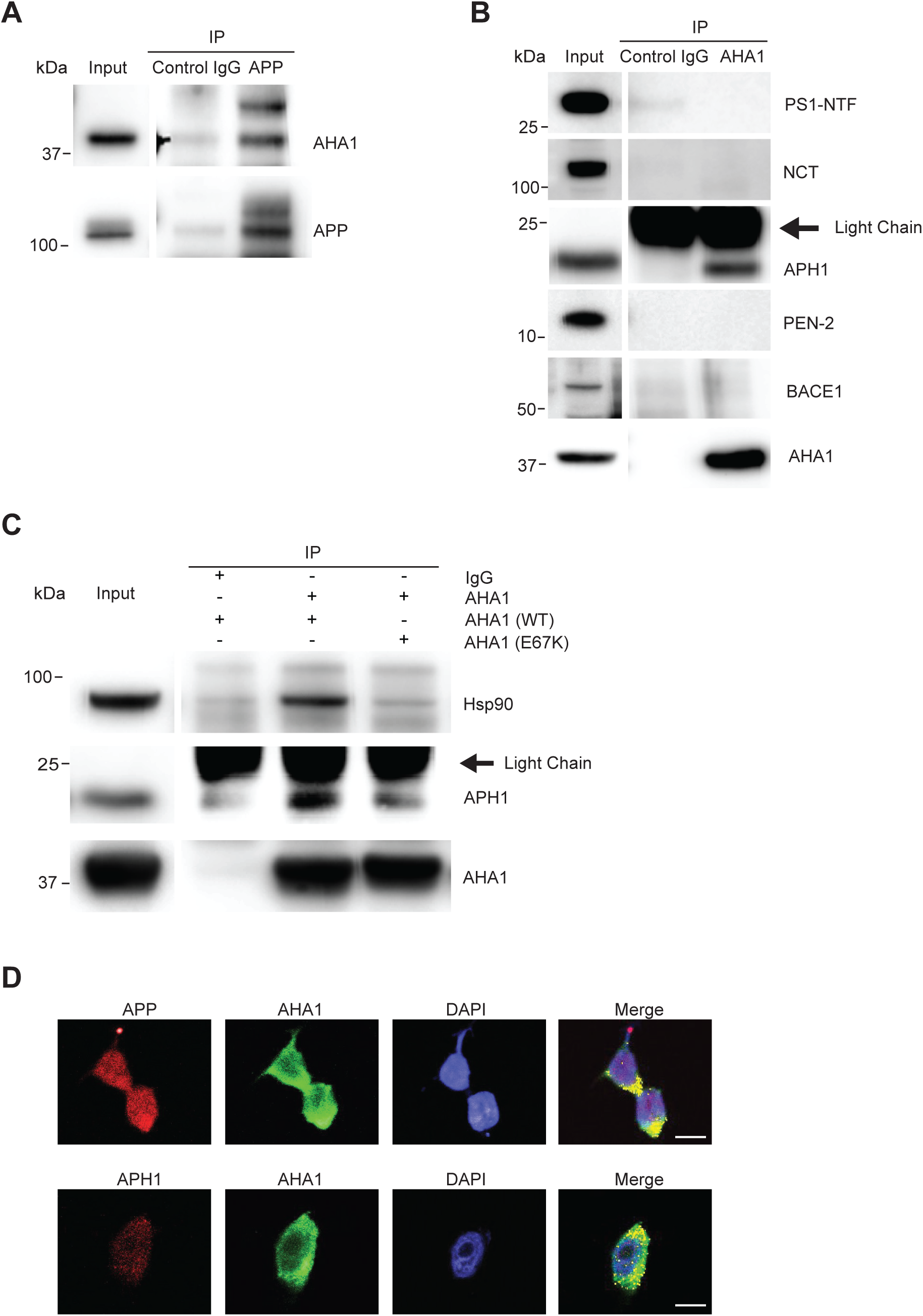
AHA1 is bound to both APP and APH1. **(A)** Lysates from HEK-APP cells at 48 h were immunoprecipitated with the APP (6E10) antibody. The obtained samples were subjected to Western blotting and analyzed using the AHA1 antibody. **(B)** Lysates from HEK-APP cells at 48 h were immunoprecipitated with the AHA1 antibody. The obtained samples were subjected to Western blotting and analyzed using antibodies that recognize γ-secretase components and BACE1. **(C)** HEK293T cells transfected with either WT AHA1 or AHA1 E67K. Lysates from cells were immunoprecipitated with the AHA1 antibody. The obtained samples were subjected to Western blotting and analyzed using Hsp90 and APH1 antibodies. **(D)** Immunofluorescence analysis showed that AHA1 colocalized with APP or APH1. HEK-APP cells were stained with secondary antibodies labeled with Alexa Fluor 488 and Alexa Fluor 568: AHA1 (green), APP (red), and APH1 (red). Nuclear boundaries were determined with DAPI (blue). Data is obtained from three independent experiments. Scale bars (D): 10μM

### AHA1 modulates the formation of the **γ**-secretase complex

As a previous study reported that knock-down of Hsp90 significantly reduced the levels of precomplex and mature complex of γ-secretase (18), we further investigated whether the AHA1/Hsp90 complex affects the formation of the γ-secretase complex. Given that the precomplex of γ-secretase is formed by APH1 and NCT before the formation of mature γ-secretase complex (4, 38),in BN-PAGE, the band at ∼ 440 kDa represents the mature γ-secretase, while the band at ∼240 kDa represents the APH1/NCT precomplex (39). Using BN-PAGE, we found that knock-down of AHA1 decreased the levels of both precomplex (Fig. 4A, lower panel at ∼240 kDa) and mature γ-secretase complex (Fig. 4A, upper panel at ∼440 kDa) in HEK-APP cells. Similarly, we also found that knock-down of AHA1 remarkably decreased the mature γ-secretase complex levels of PS1-NTF and APH1 as well as the levels of APH1 precomplex in WT MEF cells (Fig. S3A). On the other hand, co-transfection of WT AHA1+APP significantly increased the levels of both precomplex and mature γ-secretase complex compared to EV+APP control in HEK293T cells (Fig. 4B). However, co-transfection of AHA1 E67K+APP did not upregulate the levels of precomplex and mature γ-secretase complex compared to WT AHA1+APP or EV+APP (Fig. 4B). These results indicate that AHA1 is positively correlated with the formation of the γ-secretase complex and suggest that AHA1/Hsp90 complexes are essential for regulating γ-secretase assembly. We further verified our findings by treating HEK-APP cells with ND-AHA1-46 inhibitor for 24 h. We found that the formation of the APH1/NCT precomplex and mature γ-secretase complex formation was remarkably decreased in these cells compared to DMSO (Fig. S2C), indicating that AHA1 positively regulates the γ-secretase complex formation through its interaction with Hsp90.

**Figure 4.**
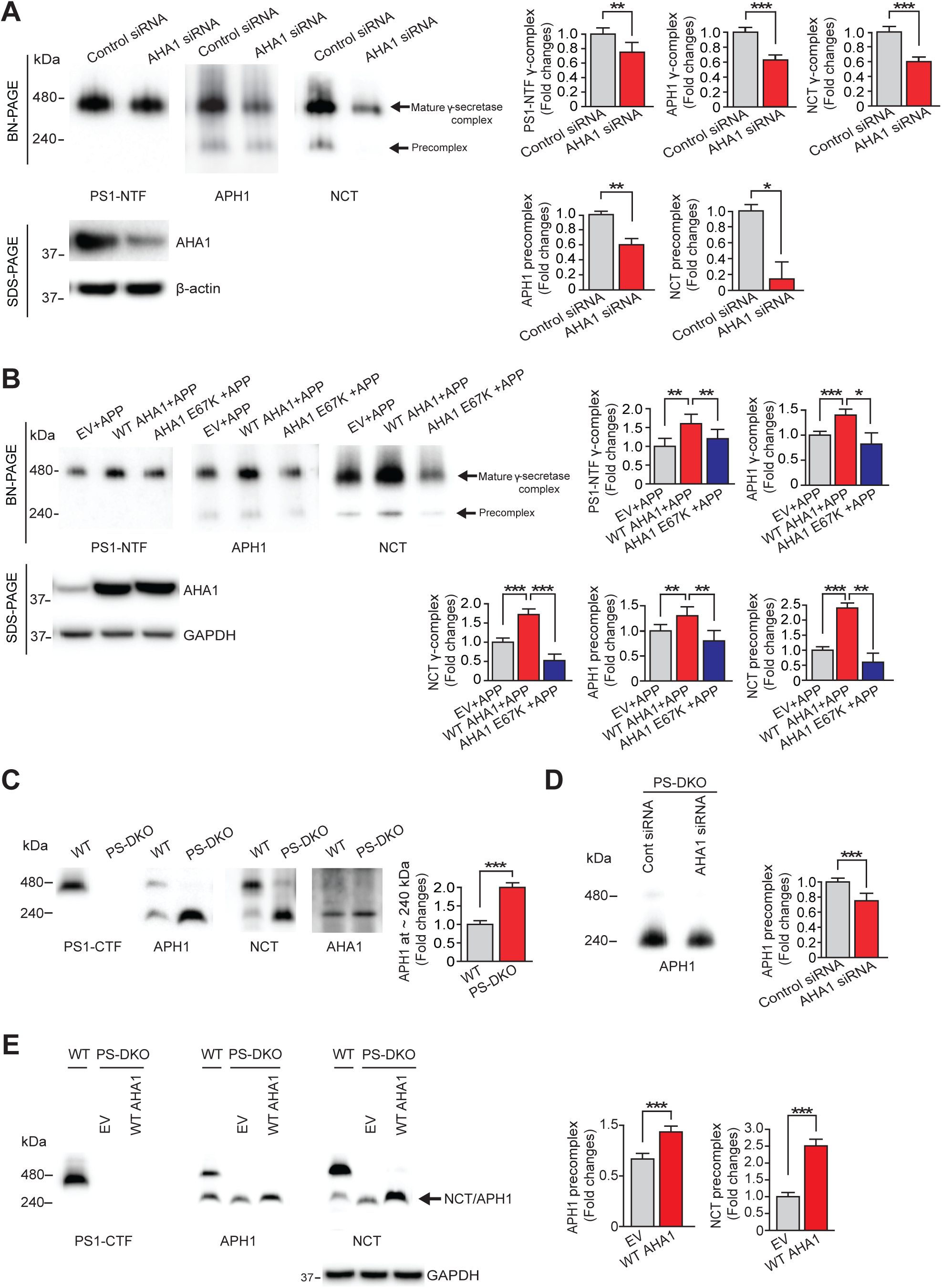
AHA1 regulates the assembly of the γ-secretase complex. **(A)** 10 μg of protein from cells transfected with control siRNA or AHA1 siRNA were subjected to BN-PAGE or SDS-PAGE and analyzed with Western blotting using the indicated antibodies. **(B)** 10 μg of protein from cells co-transfected with EV+APP or WT AHA1+APP or AHA1 E67K+APP cells were subjected to BN-PAGE or SDS-PAGE and were analyzed with Western blotting using the indicated antibodies. **(C)** 10 μg of protein from WT or PS-DKO MEF cells were subjected to BN-PAGE and analyzed with Western blotting using the indicated antibodies. **(D)** PS-DKO MEF cells transfected with control siRNA or AHA1 siRNA were subjected to BN-PAGE and analyzed with Western blotting using APH1 antibody. **(E)** PS-DKO MEF cells transfected with EV or WT AHA1 were subjected to BN-PAGE or SDS-PAGE and analyzed with Western blotting using the indicated antibodies. Data are the mean ± S.E. (error bars) from three independent experiments. *, p < 0.05; **, p < 0.01; ***, p < 0.001 as determined with Student’s *t* test and one-way ANOVA with Tukey’s multiple-comparison test.

The formation of the active γ-secretase complex is thought to begin with an intermediate APH1/NCT precomplex, which is then followed by the addition of PS and PEN-2 (4, 38). In the absence of PS, the mature γ-secretase complex cannot form, resulting in the accumulation of the APH1/NCT precomplex (39, 40). This led us to hypothesize that the AHA1/Hsp90 complex might regulate the formation of the mature γ-secretase complex by controlling the formation of APH1/NCT precomplex. We observed that in the absence of PS, mature γ-secretase complex formation was completely abolished, leading to the accumulation of APH1/NCT precomplex (∼240 kDa band) in PS-DKO MEF cells, as reported previously (Fig. 4C). Interestingly, we found that AHA1 did not incorporate into the mature γ-secretase complex but instead co-migrated with the APH1/NCT precomplex at ∼240 kDa (Fig. 4C). Additionally, we observed that the expression levels of AHA1 were elevated in PS-DKO cells compared to WT MEF cells (Fig. 4C). These findings indicate that AHA1 may play a role in promoting the assembly of the APH1/NCT precomplex, thereby facilitating the formation of the γ-secretase holo-complex through its interaction with APH1. We further investigated the role of AHA1 on γ-secretase precomplex formation in PS-DKO fibroblasts by BN-PAGE. We observed that knockdown of AHA1 significantly decreased the levels of APH1/NCT precomplex formation at ∼240 kDa in PS-DKO fibroblasts compared to control siRNA (Fig. 4D). Similar results were obtained when PS-DKO cells were treated with ND-AHA1-46 (Fig. S2C). Overexpression of WT AHA1 remarkably increased the levels of APH1/NCT precomplex formation in PS-DKO fibroblasts compared to EV control (Fig. 4E). Collectively, these findings suggest that AHA1 can modulate γ-secretase complex formation by regulating APH1/NCT γ-secretase precomplex formation, and this modulation of γ-secretase complex is positively correlated with AHA1 expression.

### AHA1 regulates abnormal A**β** production and **γ**-secretase complex formation in HEK293T cells and iPSC-derived neurons carrying FAD-linked APP mutants

Because Hsp90 stabilizes mutant proteins and prevents their degradation by interacting with them (41-43), we hypothesized that APP mutants could be a client protein of Hsp90 that is modulated by AHA1. To test this, we transiently transfected FAD-linked APP C99-I45F mutant or APP C99 as a control in HEK293T cells. Interestingly we found that overexpression of APP C99-I45F mutant remarkably increased the levels of AHA1, Hsp90, and γ-secretase molecules APH1 expression but not the expression of PS1-CTF compared to APP C99 control (Fig. 5A). Moreover, APP-V717I mutants in age- and sex-matched iPSC-derived neurons increased the Aβ42/Aβ40 ratio (Fig. S4A) and levels of AHA1, Hsp90, APP, and PS1-NTF compared to WT controls (Fig. S4B). We also found that the APP-V717I mutant increased the PS1-NTF mature γ-secretase complex formation compared to the WT control (Fig. S4C). These results suggest that FAD-linked APP mutant-mediated upregulation of AHA1 regulates γ-secretase. Furthermore, transfection of APP C99-I45F mutant did not alter the levels of C99 (Fig. 5A), suggesting that upregulation of AHA1 and Hsp90 depends on the APP C99-I45F mutation regardless of the levels of the C99 proteins. We further conducted a deeper investigation into the impact of the APP C99-I45F mutant on Aβ secretion. We found that the levels of Aβ40 secretion were dramatically reduced, and the levels of Aβ42 secretion were dramatically increased in APP C99-I45F transfected cells compared to APP C99 control. In consequence, the levels of Aβ42/Aβ40 ratio were dramatically upregulated in APP C99-I45F transfected cells (Fig. 5B) as reported previously (44). These results indicate that FAD-linked APP mutation enhanced expression of AHA1/Hsp90, which in turn contributed to the upregulation of γ-secretase levels or activity and consequently increased the Aβ42/Aβ40 ratio levels.

**Figure 5.**
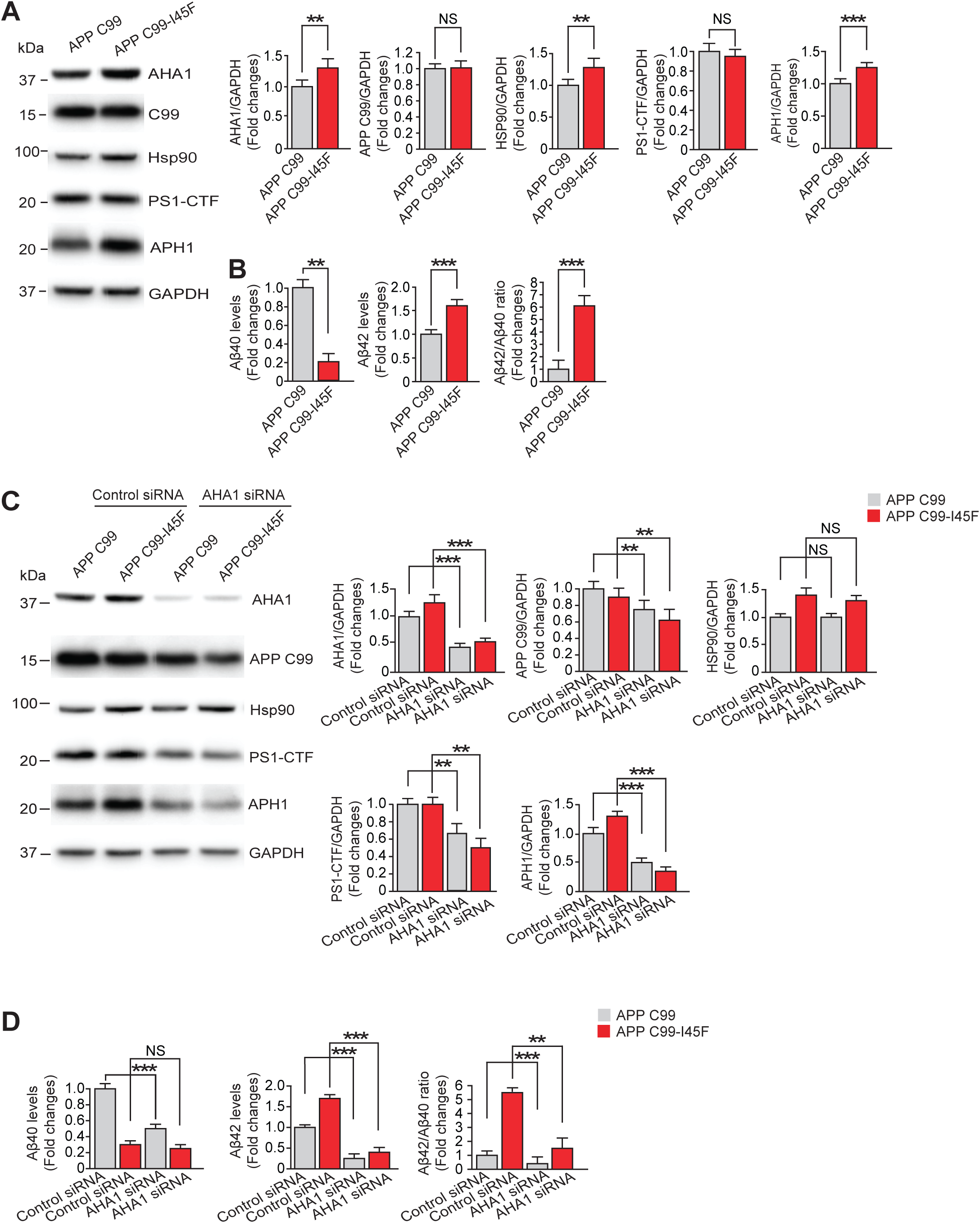
AHA1 controls abnormal. **A**β **production and** γ**-secretase complex formation in HEK-293T cells and iPSC-derived neurons carrying FAD-associated APP mutations. (A)** Transient transfection of APP C99 or FAD-linked APP C99 mutant I45F (APP C99-I45F) in HEK293T cells. The expression levels of AHA1, APP C99, Hsp90, PS1-CTF, and APH1 were detected with Western blotting. The amount of secreted Aβ40 and Aβ42 levels in the media of HEK293T cells transfected with APP C99 or APP C99-I45F were measured with an Aβ ELISA kit. Expression levels of AHA1, APP C99, Hsp90, PS1-CTF, and APH1 were assessed by Western blotting in HEK293T cells after co-expressing AHA1 knockdown and APP C99 or APP C99-I45F. **(D)** The amount of secreted Aβ40 and Aβ42 levels in the media of the HEK293T cells after co-expressing AHA1 knockdown and APP C99 or APP C99-I45F were measured with an Aβ ELISA kit. Data are the mean ± S.E. (error bars) from three independent experiments. *, p < 0.05; **, p < 0.01; ***, p < 0.001 as determined with Student’s *t* test and one-way ANOVA with Tukey’s multiple-comparison test.

We examined the effect of AHA1 knockdown on the levels of C99, Hsp90, PS1-CTF, and APH1 in HEK293T cells transfected with both C99 and C99-I45F mutants. We observed that the expression levels of C99, PS1-CTF, and APH1 were significantly decreased in AHA1 knockdown cells as compared to control siRNA of APP C99 and APP C99-I45F (Fig. 5C). However, Hsp90 did not alter after knockdown of AHA1 in these cells (Fig. 5C). To investigate whether AHA1 upregulation by the influence of FAD-linked APP mutants is essential for abnormal Aβ production, we performed a knockdown of AHA1 using siRNA in HEK293T cells transfected with both APP C99 and APP C99-I45F mutants. We found that the knockdown of AHA1 remarkably decreased the levels of Aβ40 in APP-C99 transfected cells compared to the APP C99 control siRNA (Fig. 5D). However, the knockdown of AHA1 did not significantly alter the levels of Aβ40 in APP-C99-I45F transfected cells compared to the APP-C99-I45F control siRNA (Fig. 5D). On the other hand, knockdown of AHA1 remarkably decreased the levels of Aβ42 both in APP C99 and APP-C99-I45F transfected cells compared to the control siRNA of APP-C99 and APP C99-I45F, resulting in the reduction of Aβ42/Aβ40 ratio (Fig. 5D). Taken together, these results suggest that FAD-linked APP mutant (APP C99-I45F) mutation increases the AHA1 level, which in turn contributes to an increased level of γ-secretase, mainly APH1, and leads to abnormal Aβ production.

### AHA1 regulates abnormal A**β** production and APP expression in FAD-linked PS1 mutants

Previous studies have shown that certain PS1 mutations linked to FAD upregulate the expression of APP and may produce more abnormal Aβ production. These findings were reported using various cell lines, including iPSC-derived neurons (45-49), Neuro2a (N2a) cells (50), MEF cells (51), and transgenic mice brains (52). However, the specific molecular mechanism behind this phenomenon is not well understood. Because PS1 mutations induce significant cellular stress (53), Hsp90, with its co-chaperone AHA1, plays a critical role in responding to this stress. We hypothesized that PS1 mutations might interact with Hsp90 as a client protein and that AHA1 could modulate this interaction. To test this, we used Chinese hamster ovary (CHO) cells stably overexpressing all human γ-secretase components with FAD-linked PS1 mutants (PS1-ΔE9, PS1-L166P, and PS1-L286V). We also included non-transfected (NT) parental CHO cells as a control. We found that the PS1-L286V mutant increased the expression of endogenous APP, AHA1, and Hsp90 compared to WT (Fig. 6A). However, the other PS1 mutants, ΔE9 and L166P, did not alter the expression of APP and AHA1 compared to WT. Still, they did increase the expression of Hsp90 (Fig. 6A). To gain further insight into whether increased endogenous APP expression in the PS1-L286V mutant leads to more abnormal Aβ production than WT, we transiently transfected APP695 in WT and FAD-linked PS1 mutants. We found that the PS1-L286V mutant increased Aβ42 production three-fold and increased the ratio of Aβ42/Aβ40 compared to WT (Fig. 6B), consistent with previous reports (54). On the other hand, PS1-ΔE9 and PS1-L166P mutations did not produce higher abnormal Aβ than WT. Still, the Aβ42/Aβ40 ratio increased compared to WT (Fig. 6B). Additionally, we transfected all human γ-secretase components, with either WT or FAD-linked PS1 mutant E280A, in HEK293T cells. We found that the PS1-E280A mutant increased AHA1 and endogenous APP expression compared to WT (Fig. S5A). These findings suggest that FAD-linked PS1 mutation can increase the expression of APP, AHA1, and Hsp90.

**Figure 6.**
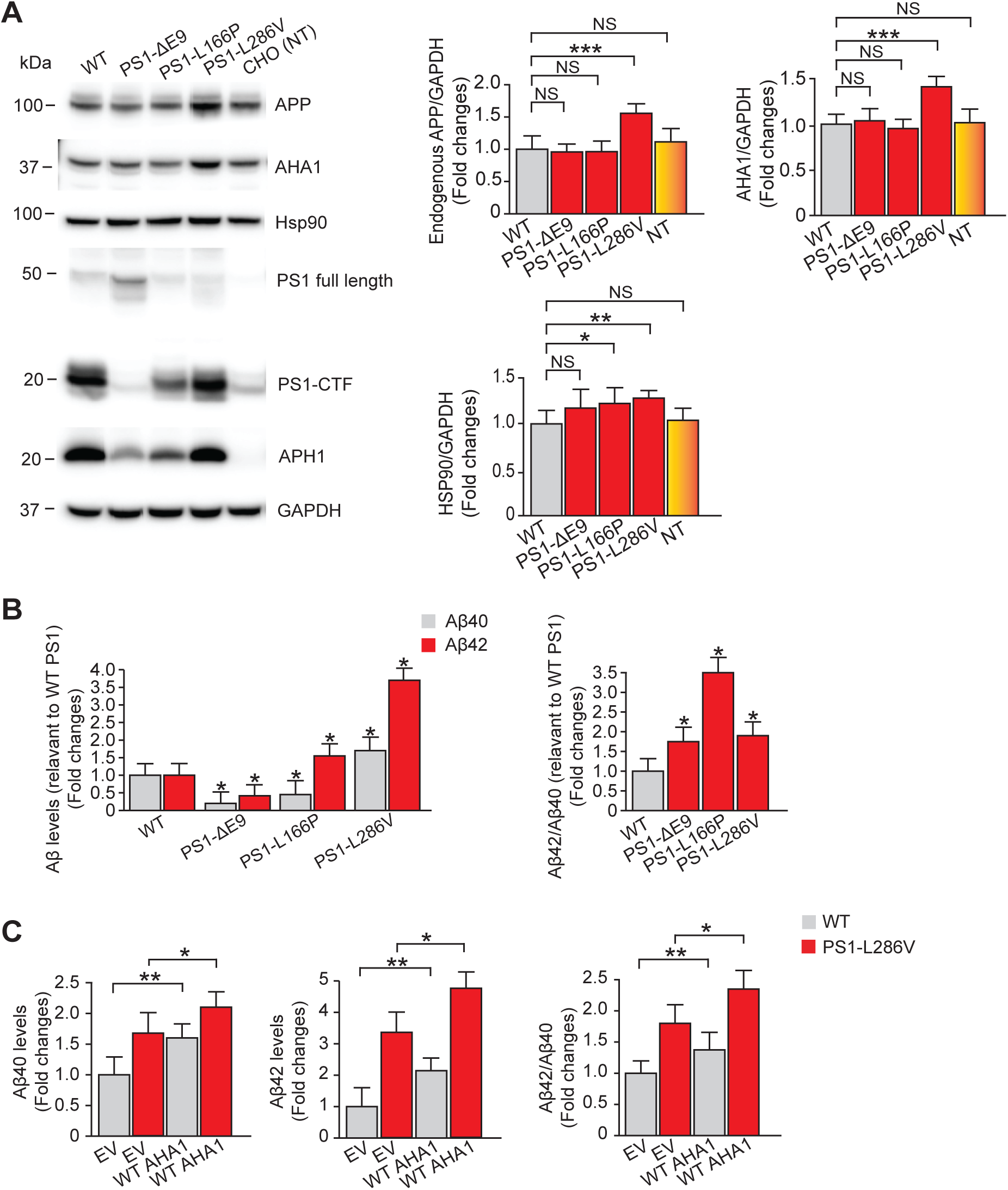
AHA1 modulates abnormal. **A**β **production and APP expression in cells carrying FAD-linked PS1 mutations. (A)** The expression levels of endogenous full-length APP, AHA1, Hsp90, full-length PS1, PS1-CTF, and APH1 were assessed by Western blotting in CHO cells stably overexpressing WT or PS1 mutants. NT-non-transfected parental CHO cells **(B)** The amount of secreted Aβ40 and Aβ42 levels in the media of CHO cells stably overexpressing WT or PS1 mutants transfected with APP695 were measured with an Aβ ELISA kit. **(C)** The levels of secreted Aβ40 and Aβ42 in the media of CHO cells stably overexpressing WT or PS1-L286V were co-transfected with APP695 and either WT AHA1 or EV control. Data are the mean ± S.E. (error bars) from three independent experiments. *, p < 0.05; **, p < 0.01; ***, p < 0.001 as determined with Student’s *t* test and one-way ANOVA with Tukey’s multiple-comparison test.

To investigate whether the increase in AHA1 expression caused by the FAD-linked PS1 mutant L286V is necessary for the abnormal production of Aβ, we conducted a co-transfection experiment of APP with either WT AHA1 or EV in CHO cells that stably express both WT and the PS1-L286V mutant. Our results showed that overexpressing AHA1 in both the WT and PS1-L286V mutant cells led to a greater increase in the production of Aβ42 compared to Aβ40, in comparison to EV (Fig. 6C). This led to a higher Aβ42/Aβ40 ratio compared to EV (Fig. 6C). Taken together, these findings suggest that AHA1 regulates the expression levels of APP and leads to increased production of abnormal Aβ in FAD-linked PS1 mutation.

## Materials and Methods

### Cell culture and DNA transfection

HEK-APP695 cells are described previously(55).CHO cells stably overexpressing all human γ-secretase components WT or PS1 mutants were previously prepared in our lab(56). HEK293T, WT, PS-DKO MEF cells were cultured in Dulbecco’s modified Eagle’s medium (DMEM) at 37 °C and 5% CO_2_. CHO cells were grown under DMEM/F12 medium at 37 °C, 5% CO_2_. All cell lines were grown in this medium supplemented with 10% fetal calf serum and 1% penicillin/streptomycin (Thermo Fisher Scientific). Transfections of DNA were performed using the Lipofectamine 3000 reagent (Thermo Fisher Scientific) according to the manufacturer’s instructions, and transfected cells were harvested at 48 h (overexpression) or 48 h (knockdown). ND-AHA1-46 inhibitor was added for 24 h in HEK-APP or PS 1/2 DKO MEF cells before harvest at 10 μM concentration.

### iPS cell culture

iPSCs were purchased from WiCell. iPSCs were differentiated into neural progenitor cells (NPCs) using STEMDiff Neural Induction Medium (NIM). iPSCs were placed into a single-cell suspension in NIM with SMADi/ROCKi (SMAD inhibitor and ROCK Inhibitor) in an AggreWell 800 plate. Embryoid bodies were cultured in the AggreWell plate for five days with NIM/SMADi partial medium changes daily. Embryoid bodies were then plated onto Matrigel-coated plates and fed daily with NIM/SMADi medium until day 12 to allow neural rosette formation. Neural rosettes were selected using Neural Rosette Selection Reagent (StemCell Tech) and plated onto Matrigel-coated dishes with NIM/SMADi. Media was changed daily for seven days, after which neural progenitor cells (NPC) were cryopreserved and split into defined Neural Progenitor Medium (StemCell Tech). NPCs were plated onto PLO/Laminin coated dishes in neural progenitor medium. The following day, the media was changed to StemDiff Forebrain Neural Differentiation Medium (StemCell Tech). Media was changed daily for seven days(57, 58). Following this, cells were plated onto PLO/laminin-coated dishes in defined Brain Phys Medium (with N2A, SM1, BDNF, GDNF, cAMP, and ascorbic acid) for neuronal maturation. Neurons were matured for 7-10 days and used for downstream experiments.

### Plasmids construct

The pcDNA plasmids encoding full-length human APP695 and APP-C99 (C-terminal 99 amino acid residues of APP) were kindly provided by Dr. L. Liu (Harvard Medical School/Brigham and Women’s Hospital, USA). The mutant form of the FAD-linked APP-C99-I45F construct and plasmids overexpressing all human γ-secretase components, including WT and FAD-linked PS1-E280A, were generated in our laboratory. The WT AHA1 and AHA1 E67K constructs were provided by Dr. Laura J. Blair (University of South Florida) and were generated using the pCMV6 backbone.

### Quantification of A**β**40 and A**β**42 levels by ELISA

Levels of secreted Aβ40 and Aβ42 in the medium were measured with sandwich enzyme-linked immunosorbent assay (ELISA) kits from Thermo Fisher Scientific according to the manufacturer’s instructions.

### AHA1 knockdown

AHA1 knockdown experiments were performed using predesigned Stealth™ siRNA against AHA1 (AHA1 siRNA) and Stealth siRNA negative control (Thermo Fisher Scientific), as previously described(59). The siRNA sequences for human AHA1: sense: 5′-CCAUGCUCCUCCAACAUUATT-3′ anti-sense: 5′-UAAUGUUGCAGGAGCAUGGGT-3′. The siRNA sequences for mouse AHA1 sense: 5′-GAACUGGACAGGUACCUTT-3′ anti-sense: 5′-AGAGGUACCUGUCCAGUUCAG-3′. Briefly, the cells were transiently transfected with 25 pmol AHA1 siRNA or control siRNA using Lipofectamine RNAiMAX (Thermo Fisher Scientific) according to the manufacturer’s instructions. The knockdown effects were examined after 48 h of incubation. The cultures were then processed for ELISA and western blot analysis.

### γ-Secretase activity assay

γ-Secretase activity was measured using a previously described γ-secretase-mediated peptide cleavage assay(60). HEK293T cells were knockdown of AHA1 by siRNA or transfected with EV or WT AHA1 or AHA1 E67K mutant grown in 150-mm culture dishes, collected in ice-cold PBS, and pelleted at 5000 rpm for 5 min. The pellet was homogenized in 500 μL of Buffer B (20 mM HEPES, pH 7.5, 150 mM KCl, 2 mM EGTA, protease, and phosphatase inhibitors) using a 27-gauge needle. The resulting homogenate was cleared at 45,000 rpm at 4 °C for 1 h. The resulting supernatant was collected, and the pellet was washed with 500 μL Buffer B and passed through a 27-gauge needle on ice. The suspension was cleared again at 45,000 rpm for 1 h at 4 °C. Supernatant was discarded, and the pellet was resuspended in 75 μL Buffer B + 1% CHAPSO and passed through a 27-gauge needle on ice. The resulting membrane samples were solubilized on a rotator at 4 °C for 2 h. Solubilized samples were cleared at 45,000 rpm for 1 h at 4 °C; supernatant (total cell membrane) was collected, and the pellet was discarded. 100 μg of protein from cells was used for γ-secretase assay. 8 μM γ-secretase fluorogenic substrate (Sigma–Aldrich) was added into samples with or without L-685,458 (Sigma–Aldrich), and the samples were incubated in γ-secretase assay buffer (100 mM Tris-HCl, pH 6.8, 4 mM EDTA, 0.5% CHAPSO) for 2 h. The values were measured by a plate reader (BioTek) at an excitation wavelength of 355 nm and an emission wavelength of 440 nm.

### Western blot analysis and Antibodies

Cells were washed with PBS and lysed in 1% digitonin or RIPA buffer (25 mm Tris-HCl (pH 7.6), 150 nm NaCl, 1% Nonidet P-40, 1% sodium deoxycholate, 0.1% SDS) with a protease inhibitor mixture (Thermo Fisher Scientific). Equal amounts of protein from the cell lysate were separated with SDS-PAGE in a 4–12% gel and blotted onto polyvinylidene difluoride membranes (Sigma–Aldrich). The membranes were incubated with primary antibodies overnight at 4 °C. Appropriate peroxidase-conjugated secondary antibodies were applied, and the membranes were visualized with Super Signal Chemiluminescence (Thermo Fisher Scientific). Membranes were stripped and reprobed with β-actin or GAPDH antibodies to normalize the loading amounts. The AHA1 antibody was purchased from Abcam. The Hsp90, BACE1, NCT, PEN2, PS1-CTF, Cleaved Notch1, β-actin, and GAPDH antibodies were purchased from Cell Signaling Technology. The APP (6E10), APH1, and PS1-NTF antibodies were purchased from BioLegend. The APP (22C11) antibody was purchased from Millipore-Sigma. The AHA1 antibody was purchased from StressMarq for immunoprecipitation. The Hsp90α antibody was purchased from Enzo Life Science for immunoprecipitation.

### Blue native–polyacrylamide gel electrophoresis (BN-PAGE)

BN-PAGE was performed as described previously(61). Cells were lysed in a native sample buffer (Thermo Fisher Scientific) containing 1% digitonin and a protease inhibitor mixture. After centrifugation at 20,000 X g at 4 °C for 30 min, the supernatant was separated on a 3–12% BisTris gel (Thermo Fisher Scientific) according to the instructions of the Novex BisTris gel system (Thermo Fisher Scientific). The transferred blot was incubated in 20 mL of 8% acetic acid for 15 minutes to fix the proteins, rinsed with deionized water, and analyzed with Western blotting.

### Immunoprecipitation (IP) Assay

HEK-APP and HEK293T cells were lysed with lysis buffer containing 1% digitonin or RIPA with a protease inhibitor mixture (Roche Applied Science), followed by centrifugation at 20,000 × g for 30 min at 4 °C. All IP steps were performed at 4 °C. Cell lysates were immunoprecipitated overnight with either anti-APP or anti-AHA1 and control IgG (Santa Cruz Biotechnology) in the presence of Protein G-Sepharose or Protein G-Agarose resin (Thermo Fisher Scientific). The beads were washed five times with lysis buffer. The samples were subjected to 4–12% gradient SDS-PAGE and transferred to a polyvinylidene difluoride membrane for Western blotting analysis.

### Immunostaining

HEK-APP cells were fixed in 2% paraformaldehyde. Next, they were permeabilized with 0.1% Triton X-100 and blocked for 45 min in 10% normal goat serum in PBS. Fixed cells were incubated with primary antibodies (AHA1 and APP or APH1) for 12 h at 4 °C. Immunofluorescent labeling was carried out with Alexa Fluor 488– and Alexa Fluor 568–tagged secondary antibodies (Thermo Fisher Scientific). The mounting was done with Vectashield reagent with DAPI (Vector Labs.). Images were captured on a confocal microscope (Leica) using an oil-immersion plan Apo ×60 A/1.40 numerical aperture objective lens.

### Statistical analysis

Statistical analysis was performed using a statistical package, GraphPad Prism 10 software (GraphPad Software, San Diego, CA). All values are presented as the mean ± S.E. of at least three independent experiments. Student’s t-test was used to determine whether the results were significantly different between the two groups. We compared group differences with a one-way analysis of variance (ANOVA) followed by Tukey’s multiple-comparison test for three or more groups against a control group. A p-value of <0.05 was considered to represent a significant difference.

## Discussion

In this study, we found that the Hsp90 co-chaperone AHA1 exerts a substantial influence on the production of Aβ. Our data indicates that AHA1 plays a crucial role in regulating the levels of Aβ by interacting with both APP and APH1. Specifically, AHA1 modulates APP levels and influences the formation of the pre-complex and mature γ-secretase complex. This modulation ultimately enhances the activity of γ-secretase. Furthermore, we found that using an Hsp90-binding-defective AHA1 E67K mutant or a pharmacological inhibitor that blocks the AHA1/Hsp90 complex resulted in a reduction in both the pre-complex and mature complex of γ-secretase. These results suggest that the AHA1 can modulate γ-secretase by pathways dependent on Hsp90 interaction. However, with the AHA1 E67K mutant or ND-AHA1-46 inhibitor, the expression of APP was increased compared to EV and DMSO. These results revealed that AHA1 can modulate the APP level by pathways independent of Hsp90 interaction. Our findings are consistent with previous findings that AHA1 can function as an Hsp90-independent, autonomous chaperone. A previous study suggests that most AHA1 exists in cells as a self-contained protein independent of high-molecular-weight Hsp90 protein complexes, and AHA1 itself can prevent the aggregation of denatured rhodanese and luciferase (29). A recent study found that Dicer1 is a client protein of AHA1, and AHA1 can influence the expression of Dicer1 independently of its interaction with Hsp90 (34). Another study suggests that SULT1A1 is a client of AHA1 but not of Hsp90 (30). These findings indicate that AHA1 may have a role beyond its known function as a potent stimulator of Hsp90’s ATPase activity.

It is believed that targeting AHA1 with specific inhibitors could be a promising approach to reducing tau pathology in AD (62, 63). However, the role of AHA1 inhibitors in reducing γ-secretase assembly formation was previously unknown. Our discovery revealed that the ND-AHA1-46 inhibitor significantly reduced the formation of the γ-secretase components and its complex. Still, it increased the levels of APP. The AHA1 inhibitor decreases tau aggregation and the formation of the γ-secretase complex, showing potential for treating AD. Nevertheless, further research is needed to develop an inhibitor that can reduce tau and Aβ production without affecting the expression level of APP. It would also be important to show that other essential γ-secretase substrates such as Notch1 are not unduly affected.

APP and PS1 autosomal dominant mutations are causative of FAD (6, 64). The upregulation of Hsp90 expression is a natural response to cellular stress, which includes responding to mutations. This upregulation helps to stabilize mutant proteins, allowing cells to maintain their proper function (43, 65). In this regard, it was previously reported that cells carrying FAD-linked APP-I45F mutant increased the expression of Hsp90 (66). In this study, we used the APP C99 Iberian (I45F) mutation to show that it substantially increases Aβ42 production while decreasing Aβ40, resulting in a significant increase in the Aβ42/Aβ40 ratio, considered a crucial indicator of AD pathogenicity. Here, we also uncovered that APP-C99-I45F mutations increased AHA1, Hsp90, and APH1 levels. However, the levels of PS1-CTF remained unchanged. Knockdown of AHA1 led to a decrease in C99, PS1-CTF, and APH1 levels. As a result, there was a decrease in the Aβ42/Aβ40 ratio. Interestingly, APP C99-I45F mutation increased APH1 levels, and recent genome-wide association studies identified APH1 as an AD risk gene (67). Furthermore, an upregulation of APH1 expression was seen in AD patients compared to the control group(68). Previous studies reported that iPSC-derived neurons carrying APP-V717I mutation increased the levels of APP and tau (58). Consistent with previous observations that the APP-V717I mutation elevates APP expression, we additionally found increased levels of Hsp90, AHA1, and PS1-CTF, collectively leading to aberrant Aβ generation.

PS1 mutations have been associated with increased cellular stress, such as endoplasmic reticulum (ER) stress(69), oxidative stress(70). Lopez-Toledo *et al.* (71) found that expression of Hsp90 was significantly higher in the fibroblasts of FAD patients carrying PS1-A246E and PS1-M146L mutations. A recent study suggests that PS1-E280A mutation increases the levels of Hsp90 compared to sporadic AD in postmortem human brain samples(72). Other studies suggest that human iPSC neurons expressing the PS1-A246E mutant increase the levels of Hsp90, APP, and tau (47, 48). Moreover, iPSC-derived neurons carrying PS1-L150P and PS1-E120K mutations increase APP and tau levels compared to control iPSC (45, 46). Furthermore, transgenic PS1-M146V knock-in mice showed increased levels of APP(52). Other reports suggest that neuroblastoma N2A cells expressing PS1-P264S mutant increased APP levels (50). Despite increasing evidence, the molecular mechanism underlying the relationship between increased APP expression and FAD-linked PS1 mutations remains unexplored. In this study, we reveal for the first time that AHA1 plays a role in regulating APP expression in FAD-linked PS1 mutants. Our research indicates that the PS1-L286V mutant increases CHO cells’ endogenous APP, AHA1, and Hsp90. Furthermore, the PS1-L286V mutant increases the abnormal Aβ production compared to the WT. Cells overexpressing cells WT AHA1 resulted in higher abnormal Aβ output levels than EV. Additionally, we found that the PS1-E280A mutation increases levels of both APP and AHA1. Several FAD mutants have been discovered to increase the Aβ42/Aβ40 ratio, leading to early amyloid deposition in the brain (73-77). Therefore, further research is needed to investigate the connection between the upregulation of AHA1/Hsp90 and the abnormal production of Aβ and accumulation of tau in FAD. Future studies using AHA1 knockout or tissue-specific AHA1 deletion mice would provide valuable insight into the role of AHA1 in AD progression, although no fully characterized AHA1 null mouse line has been widely reported.

Together, our research has shown that AHA1 plays a crucial role in the production of Aβ. We have discovered that AHA1 regulates the expression of the APP and the assembly of γ-secretase. By disrupting the complex formation between Hsp90 and AHA1, we can reduce the assembly of γ-secretase. Additionally, we observed that AHA1 is up-regulated and stimulates the expression of APP or APH1 under FAD-linked mutations, indicating a molecular connection between AHA1/Hsp90 and FAD-linked APP and PS1 mutations. Our findings suggest that targeting AHA1 could be a promising approach for developing treatments that control Aβ levels, which are critical in AD. Our findings may offer significant implications for the development of new AD treatments.

## Supporting information

Supplementary Figures and Legends for Noorani et al.

## Acknowledgments

We are grateful to Dr. Laura J. Blair (University of South Florida) for her valuable suggestions and insightful comments during the preparation of this work, and for generously providing the WT AHA1 and AHA1 E67K mutant plasmid constructs. We also thank Dr. L. Liu (Harvard Medical School/Brigham and Women’s Hospital) for providing HEK-APP, HEK293T, WT, PS1/2-DKO MEF cells, and the APP-C99 construct. We thank Caitlin Overmeyer for the APP-C99-I45F construct and Dr. Tanbir Ahammad for the WT and PS1-E280A constructs. We also thank Dr. Omar Quintero-Monzon for producing CHO cells stably overexpressing all human γ-secretase components with FAD-linked PS1 mutants. Confocal microscopy was performed at the Microscopy & Analytical Imaging Laboratory at the University of Kansas.

## Author contributions

Conceptualization: A.A.N., M.S.W.; Data curation: A.A.N., S.I., K.C., and H.M.W.; Investigation: A.A.N. and S.I.; Data analysis: A.A.N., S.I., B.S.J.B., M.S.W. and K.Z.; Writing-original draft: A.A.N., S.I.; Writing-review and editing: M.S.W., B.S.J.B and K.Z.; Supervision: M.S.W.; Funding acquisition: M.S.W.; Project administration: M.S.W.

## Abbreviations—The abbreviations used are

AD: Alzheimer’s disease
FAD: Familial Alzheimer’s disease
SAD: Sporadic Alzheimer’s disease
Aβ: amyloid beta
AHA1: activator of Hsp90 ATPase homologue 1
Hsp90: heat shock protein 90
NCT: nicastrin
APH1: anterior pharynx-defective-1
APP: amyloid precursor protein
BACE1: β-site APP-cleaving enzyme 1
PS: presenilin
PS1-CTF: C-terminal fragment of presenilin 1
PS1-NTF: N-terminal fragment of presenilin 1
PEN-2: presenilin enhancer-2
NICD: notch intracellular domain
HEK: human embryonic kidney
CHO: Chinese hamster ovary
NT: non-transfected
PS-DKO MEF: presenilin 1 and 2 double knockout mouse embryonic fibroblast
BN-PAGE: blue-native polyacrylamide gel electrophoresis
IP: immunoprecipitation
EV: empty vector
BisTris: bis(2-hydroxyethyl) aminotris (hydroxymethyl) methane
ANOVA: analysis of variance

## Data availability

All data are included in the article.

