## Supplementary Figures and Legends for Noorani et al. for "AHA1 regulates Aβ production via modulation of APP expression and γ-secretase assembly"

**Running title:** AHA1 regulates γ-secretase and APP expression

### Supplementary Figure Legends

**Figure S1. ND-AHA1-46 inhibits interaction between Hsp90 and AHA1. (A)** Chemical structure of ND-AHA1-46. The 1,5-bisubstituted-1,2,3,4-tetrazole containing AHA1/Hsp90 disruptor ND-AHA1-46. **(B)** Immunoprecipitated AHA1 from HEK293T cells treated with DMSO or ND-AHA1-46 10 µM for 24 h was analyzed by Western blotting.

**Figure S2. ND-AHA1-46 reduces γ-secretase complex formation. (A)** The expression levels of full-length APP, PS1-NTF, NCT, and APH1 in HEK-APP cells after treatment of DMSO or ND-AHA1-46 10uM were detected with Western blotting. **(B)** 10 µg of protein from DMSO or ND-AHA1-46 treated cells were subjected to BN-PAGE or SDS-PAGE and analyzed with Western blotting using the indicated antibodies. **(C)** PS-DKO MEF cells treated with DMSO or ND-AHA1-46 10uM for 24 h. 10 µg of protein from DMSO or ND-AHA1-46 treated cells were subjected to BN-PAGE and analyzed with Western blotting using the APH1 antibody. Data are the mean ± S.E. (error bars) from three independent experiments **, $p < 0.01$; ***, $p < 0.001$ as determined with Student's *t* test.

**Figure S3. Knockdown of AHA1 decreases γ-secretase complex formation. (A)** 10 µg of protein from WT MEF cells after knockdown of AHA1 were subjected to BN-PAGE or SDS-PAGE and analyzed with Western blotting using the indicated antibodies. **, $p < 0.01$; ***, $p < 0.001$ as determined with Student's *t* test.

**Figure S4. APP-V717I mutation increases the levels of AHA1, Hsp90, APP, PS1-NTF, and γ-secretase complex. (A)** The amount of secreted Aβ40 and Aβ42 levels in the media of iPSC-derived neurons with WT or APP-V717I mutation were measured with an Aβ ELISA kit. **(B)** The expression levels of AHA1, Hsp90, full-length APP, and PS1-NTF in iPSC-derived neurons with WT or APP-V717I mutation were detected with Western blotting. **(C)** 10 µg of protein from WT or APP-V717I cells was subjected to BN-PAGE and analyzed with Western blotting using PS1-NTF antibody. Data are the mean ± S.E. (error bars) from three independent experiments **, $p < 0.01$; ***, $p < 0.001$ as determined with Student's *t* test.

**Figure S5. PS1-E280A mutation increases the levels of AHA1 and APP. (A)** The expression levels of AHA1, full-length APP, PS1-NTF, NCT, and PEN2 were assessed by Western blotting in HEK293T cells after transfection of overexpressing WT or PS1-E280A. Data are the mean ± S.E. (error bars) from three independent experiments. *, $p < 0.05$; **, $p < 0.01$; ***, $p < 0.001$ as determined with Student's *t* test.

**Noorani et. al.,**
**Fig. S1.**

**A**

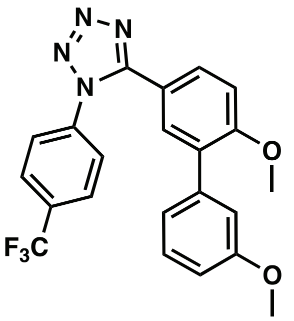


**ND-AHA-46**

**B**

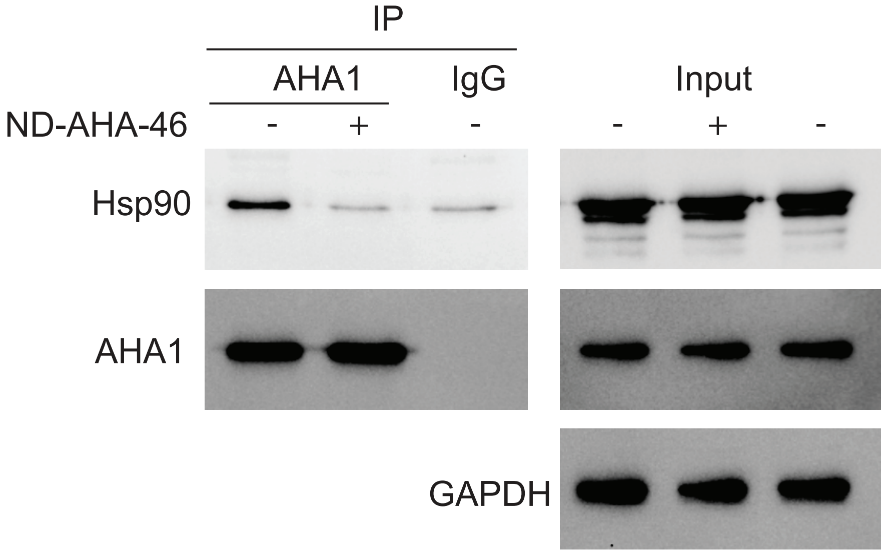

**Noorani et. al.,**
**Fig. S2.**

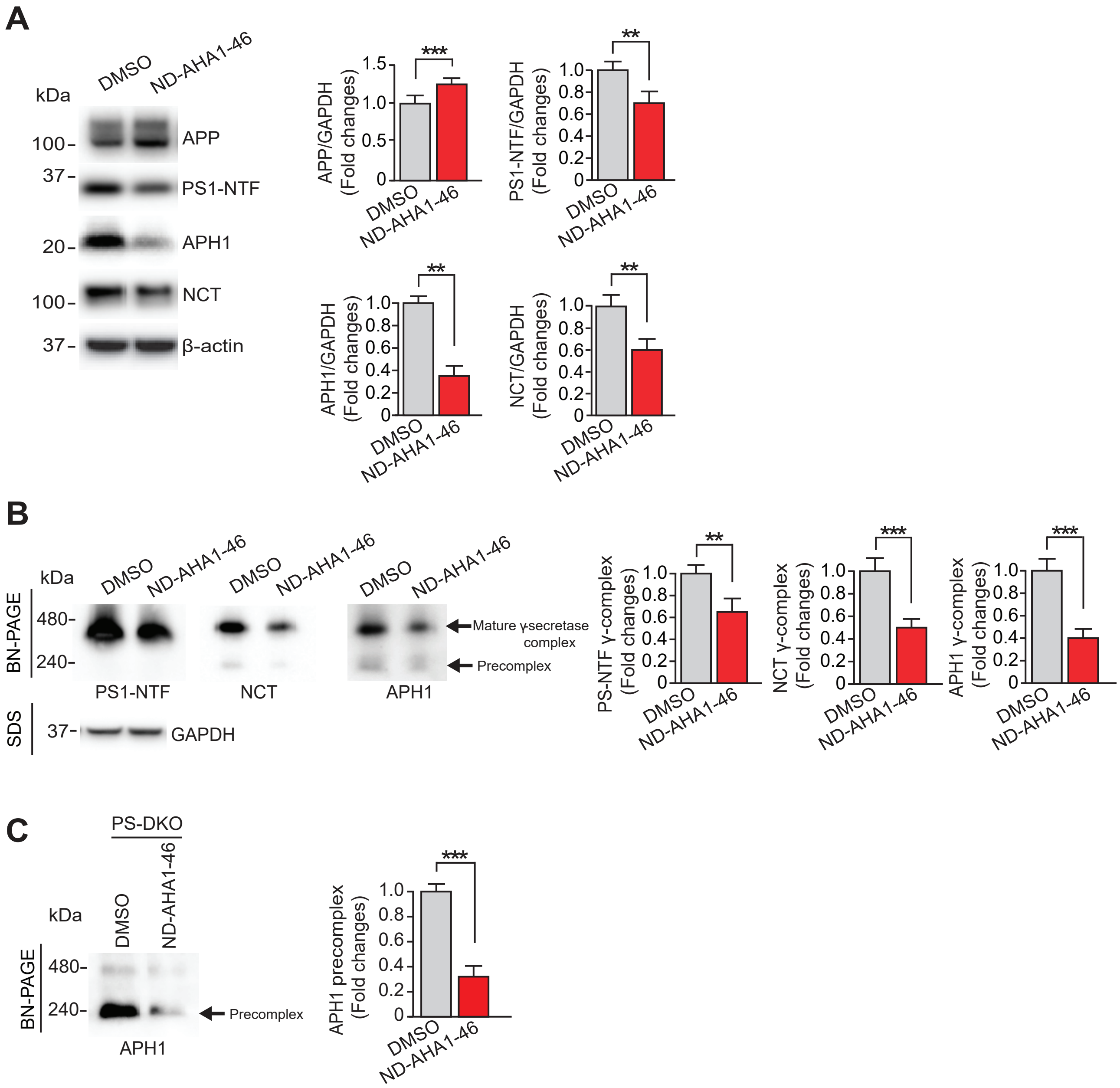

**Noorani et. al.,**
**Fig. S3.**

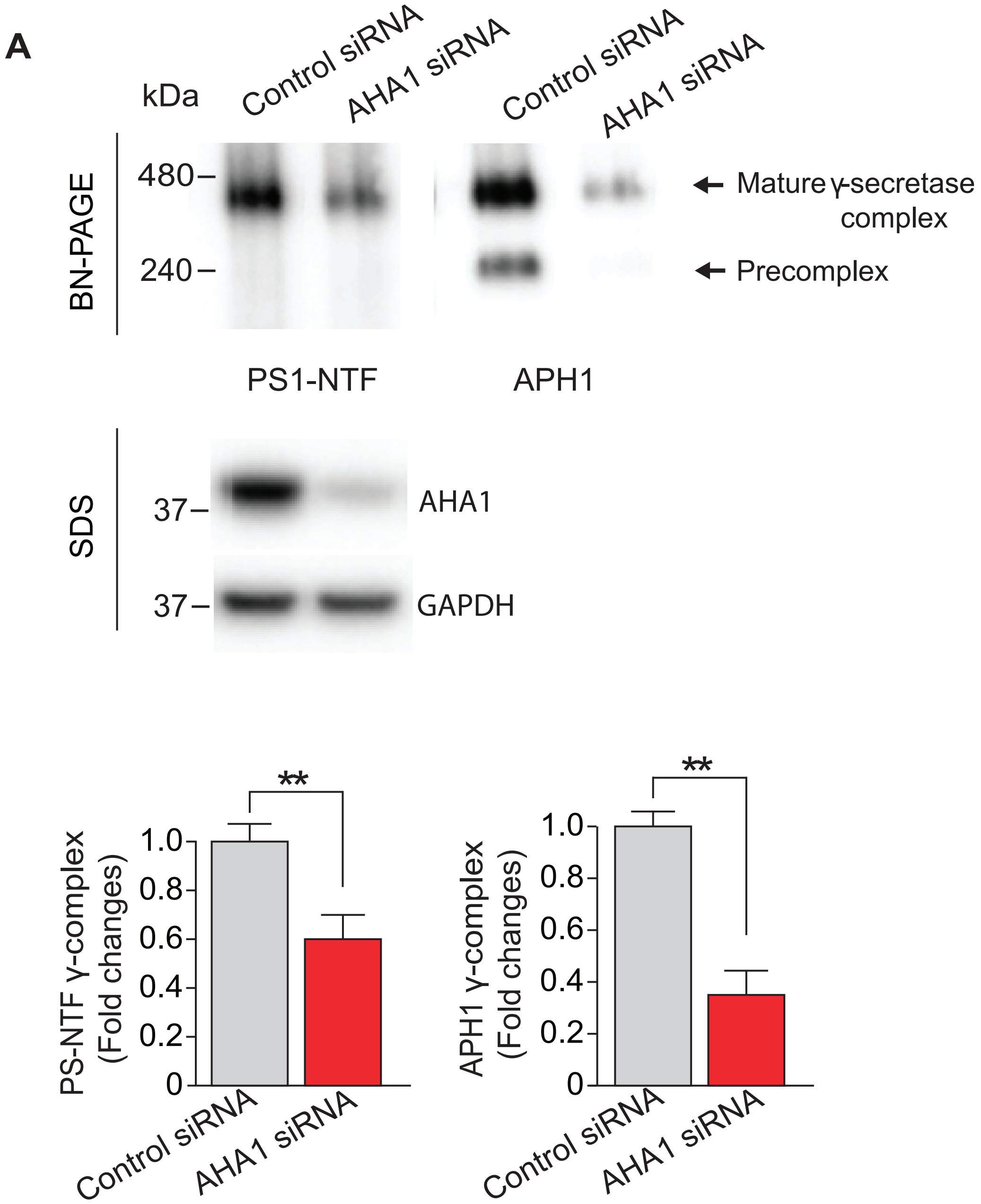

**Noorani et. al.,**
**Fig. S4.**

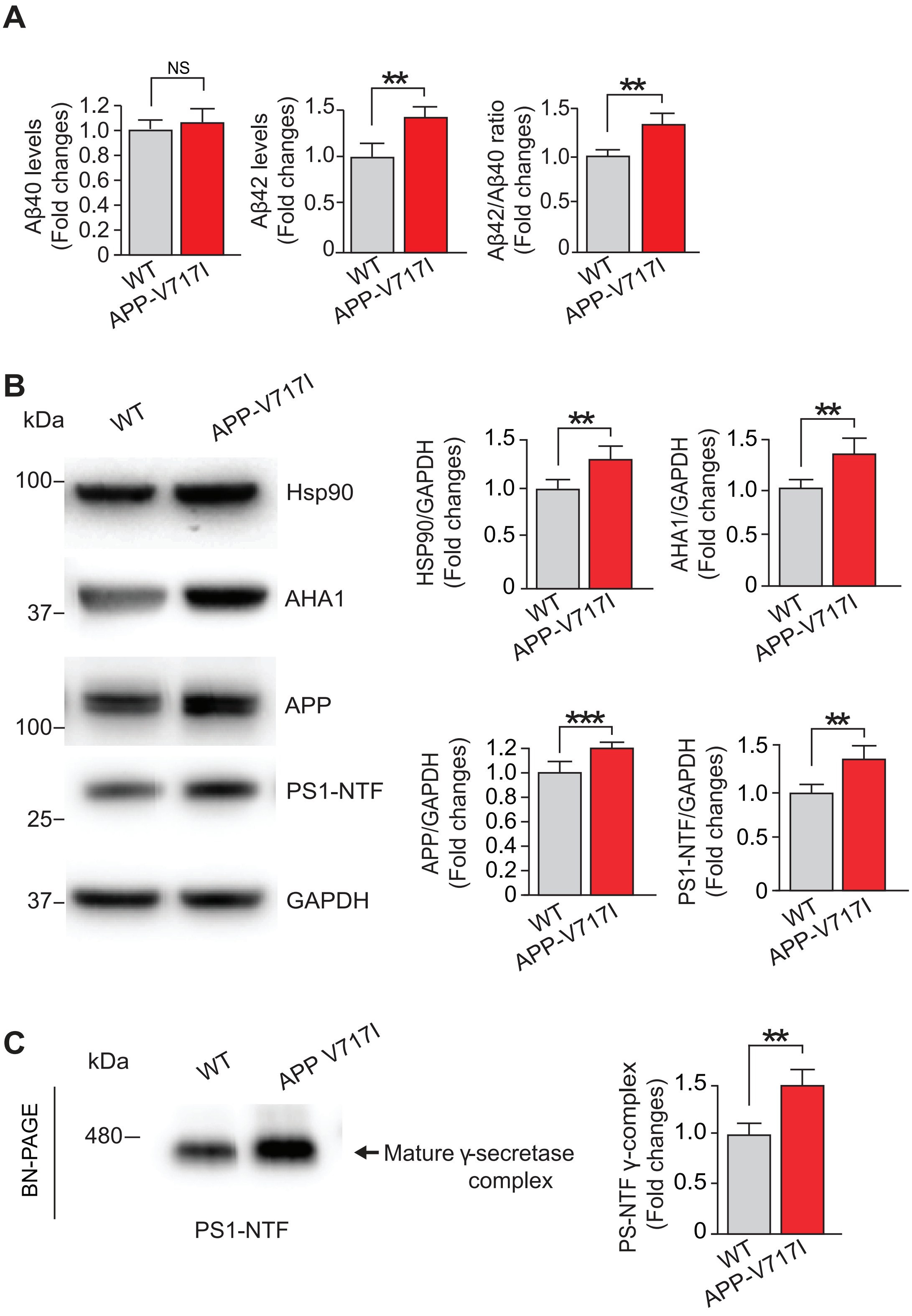

**Noorani et. al.,**
**Fig. S5.**

**A**

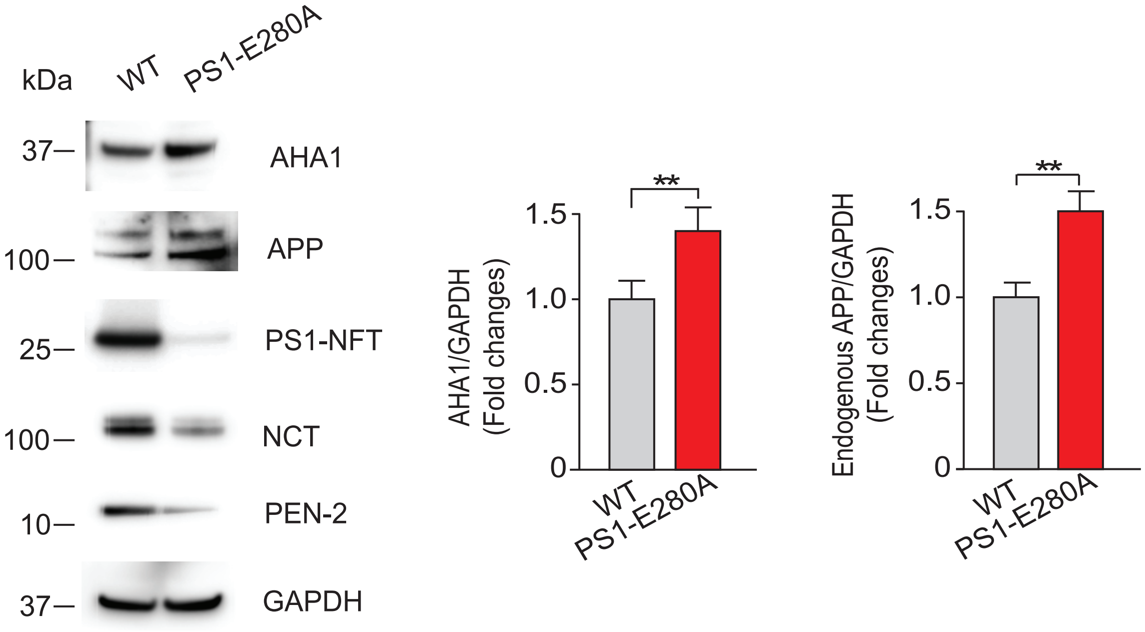
